# A hybrid reconstruction of the physical model with the deep-learning that improves structured illumination microscopy

**DOI:** 10.1101/2022.10.04.510914

**Authors:** Jianyong Wang, Junchao Fan, Bo Zhou, Xiaoshuai Huang, Liangyi Chen

## Abstract

In handling raw images with low signal-to-noise (SNR) ratios, conventional algorithms of structured illumination microscopy are prone to artifacts, while deep-learning-based (DL) algorithms may lead to degradation and hallucinations. We propose a hybrid that combines the physical inversion model with a Total Deep Variation regularization. In super-resolving from low SNR images such as actin filaments, our method outperforms conventional or DL methods in suppressing artifacts and hallucinations while maintaining resolutions.

---

Despite being the method of choice for live-cell super-resolution (SR) imaging, structured illumination microscopy (SIM) suffers from reconstruction artifacts as its image restoration confers an ill-posed inverse problem^1–3^. The problem is exacerbated under fast live-cell SIM imaging with sub-millisecond exposures or excessive photobleaching, as limited photon fluxes emitted from fluorophores yield lower SNR raw images^1,2,4–7^. Various model-based restoration methods have been developed to suppress SIM artifacts, including TV-SIM^9^, HiFi-SIM^10^, and Hessian-SIM^8^. However, these methods are designed to handle specific artifacts and may not completely resolve the problem. People have proposed end-to-end deep-learning (DL) based reconstruction methods to suppress different artifacts indiscriminately^11,12^. However, DL-based methods may suffer from hallucinations and generally reduced resolution. For example, current DL methods often incorrectly predict mitochondrial cristae structures in live cells.

By combining the physical SIM reconstruction procedure with a Total Deep Variation (TDV) regularizer^13^, we proposed a hybrid restoration method, TDV-SIM, to suppress artifacts and maintain resolution simultaneously. We transformed the SIM reconstruction into an optimization problem and constructed an objective function **[Eq. (1)]** composed of the fidelity term *D*(*f, g*) based on the physical model (**Supplementary Note 1**) and the TDV regularization term *R*(*f*) based on DL.

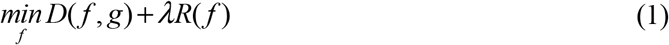

where *f* is the target image to be estimated, g is the inverse Fourier transform of the high and low-frequency information separated from the SIM raw data, and λ is the weight parameter of the regularization term. By optimizing the objective function with the gradient descent algorithm **[Eq. (2)]**, TDV-SIM can reconstruct SR-SIM images that preserve the high-frequency information more faithfully than pure DL-based methods, and suppress artifacts more effectively than pure model-based methods.

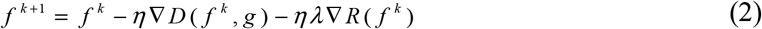

where *η* is the step size. The entire reconstruction pipeline is shown in **Fig. 1a**, where *f* ^0^ is the initial SIM image obtained by Wiener deconvolution and *f* ^*T*^ is the final reconstruction after *T* iterations. The computation pipeline of ∇*R*(*f*) is shown in **Fig. 1b**. Compared to the ground truth (GT) image of actin filaments (averages of multiple Wiener-processed images, **Fig. 1c**), we quantized the peak signal-to-noise (PSNR, **Fig. 1d, top**) and structural similarity index measure (SSIM, **Fig. 1d, bottom**) values of TDV-SIM reconstructions with different weight parameter λ and iteration number T. Through human inspection (**Fig. 1e**), artifacts may not be suppressed entirely if λ (or *T*) is too small; in contrast, if λ (or *T*) is too large with a fixed *T* of 25 (or a λ of 2.5), genuine signals may be removed incorrectly. Thus we set the optimal parameters to be 2.5 and 25 for λ and *T*, respectively.

**Fig. 1.**
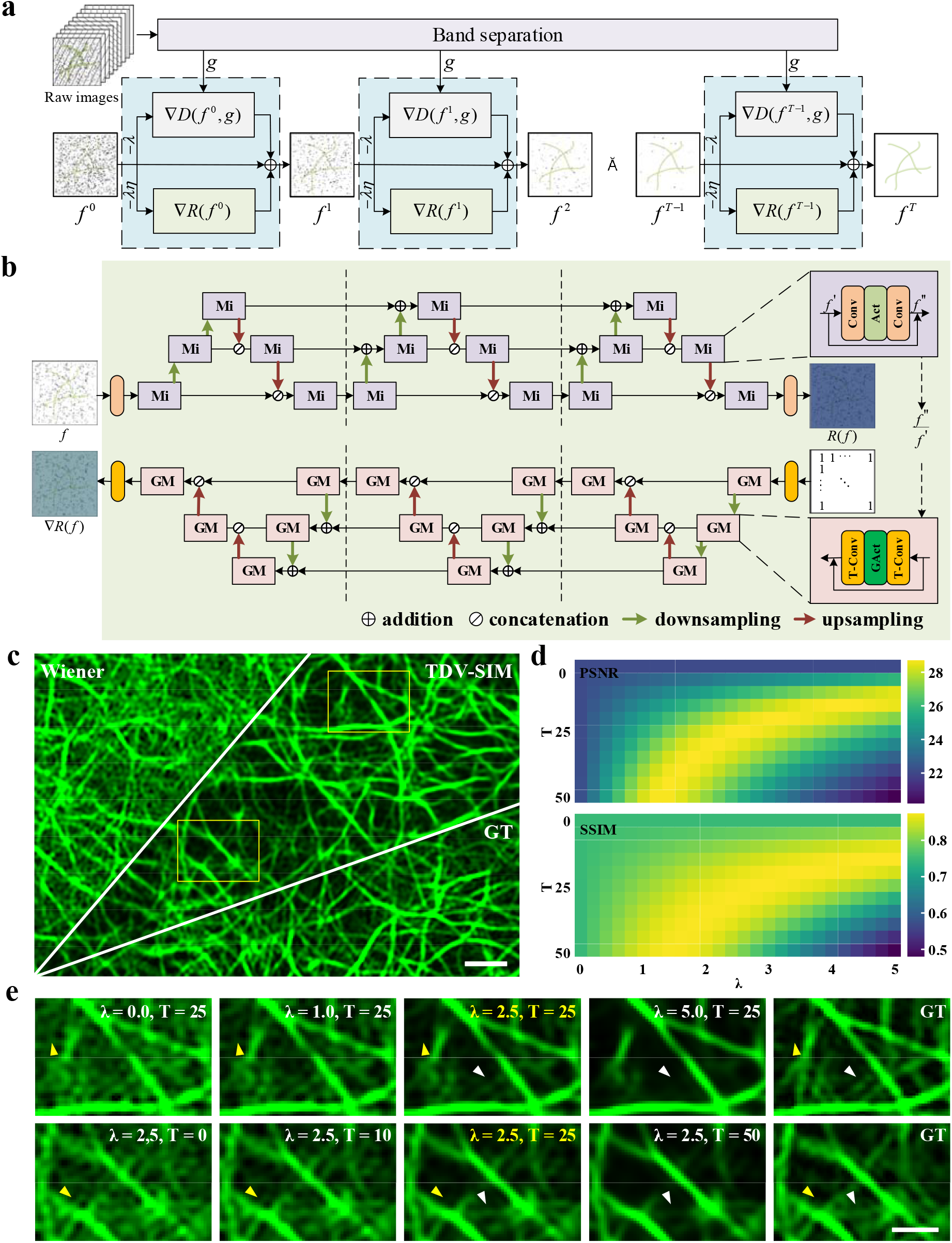
TDV-SIM diagrams and parameters selection. **(a)** TDV-SIM reconstruction pipeline. (**b**) Visualization of TDV and its gradient. **Mi** is a residual structure micro-block, and **GM** is its gradient. **Conv** is the convolution layer, and **T-Conv** is its gradient. **Act** is the activation layer, and **GAct** is its gradient. (**c**) Actin filaments SIM SR image. (**d**) PSNR and SSIM of TDV-SIM reconstructions with different *λ* and *T*. (**e**) Magnified views of the boxed regions in (**c**). Yellow arrowheads highlight artifacts not eliminated with too small *λ* or *T*. White arrowheads indicate incorrectly removed signals with too large *λ* or *T*. Scale bars: (**c**) 1 μm; (**e**) 0.5 μm.

First, we compared TDV-SIM with other reconstruction methods, including physical-model-based (Wiener deconvolution^3^, HiFi-SIM, Hessian-SIM) and pure DL-based methods(scU-Net^11^, DFCAN^12^) using synthetic images with known GT (**Fig. S1**). TDV-SIM confers balanced performance in generating SR images of high SSIM, low normalized root-mean-square error, and low artifacts among all reconstruction methods. Next, we examined dynamic actin filaments and ER in live cells observed with short exposures (actin: 1 ms, **Fig. 2a**; 2.7 ms, **Fig. S2a**; ER: 0.789 ms, **Fig. 2d**). Despite the improved reconstructions compared to the Wiener deconvolution, HiFi-SIM and Hessian-SIM still produced artifacts due to noise amplification in background regions with low SNR. TDV-SIM produced more continuous actin filaments (**Fig. S2e**) with fewer artifacts but comparable SSIM values and resolutions to the conventional reconstruction methods (**Fig. 2b,e,f-j**, and **Fig. S2d**). In contrast, pure DL-based methods led to reconstruction with fewer artifacts at the price of reduced resolution and decreased SSIM values. In addition, we often observed inaccurate inferences at the intersections of actin filaments and ER (yellow arrows in **Fig. 2c**,**e**, and **Fig. S2c**). Together with the incorrectly inferred actin filaments at regions with extremely low fluorescence intensity (**Fig. S3**), these resembled the “hallucination effects” of pure DL methods^14^, which was abolished by the TDV-SIM method.

**Fig. 2.**
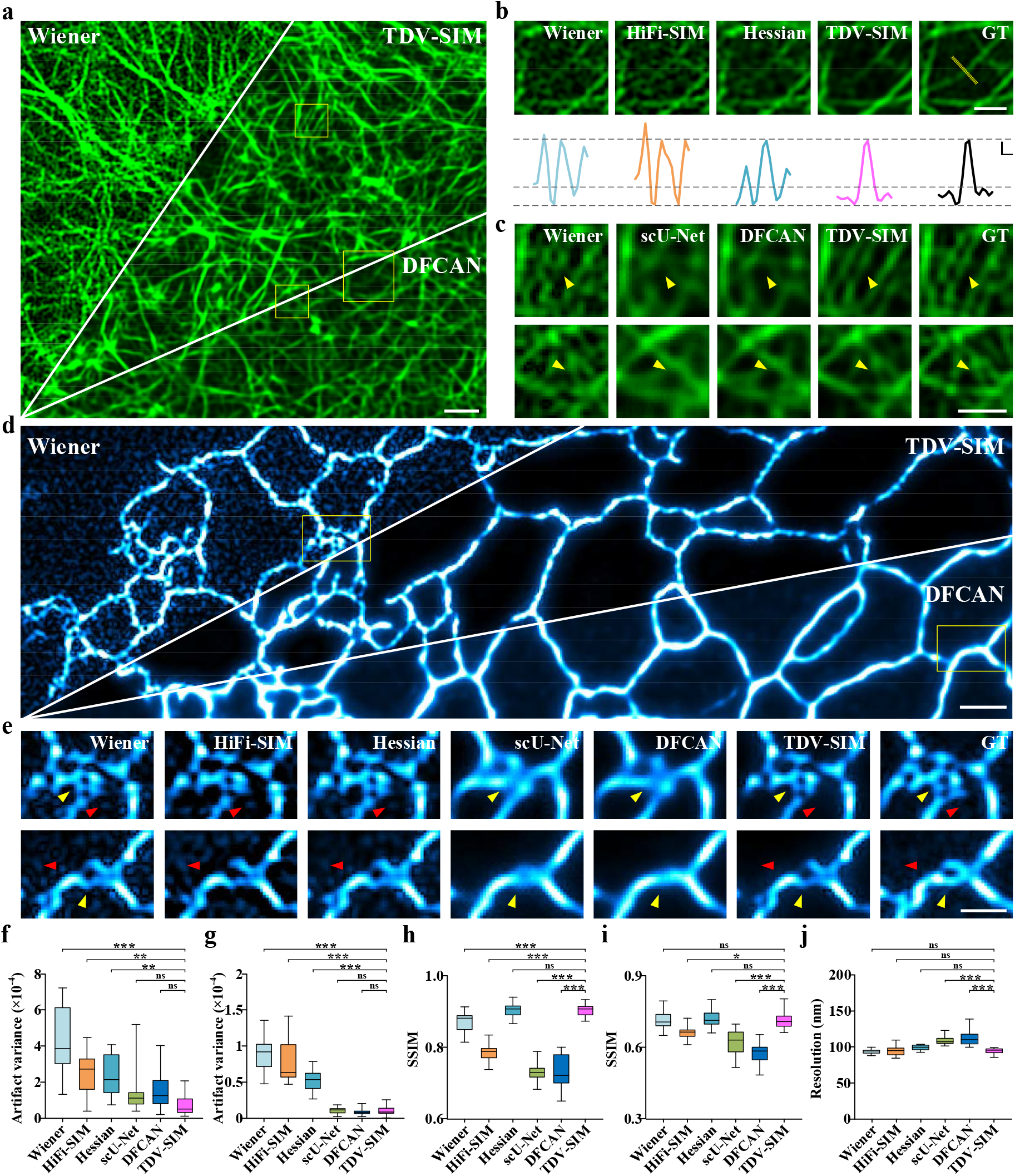
TDV-SIM outperforms other reconstruction algorithms in suppressing artifacts and hallucinations while maintaining resolution. (**a**) Actin filaments under the SR-SIM. (**b**) Magnified views of the larger boxed region in (**a**) reconstructed by Wiener deconvolution, HiFi-SIM, Hessian-SIM, and TDV-SIM. The GT image is shown as the reference. Profiles along the yellow line are on the bottom. (**c**) Magnified views of the smaller boxed regions in (**a**) reconstructed by Wiener deconvolution, scU-Net, DFCAN, and TDV-SIM. GT images are shown as references. (**d**) ER under the SR-SIM. (**e**) Magnified views of the boxed regions in (**d**) reconstructed by Wiener deconvolution, HiFi-SIM, Hessian-SIM, scU-Net, DFCAN, and TDV-SIM. The GT images are shown as references. (**f-g**) Artifact variances of actin filaments (**f**) or ER tubules (**g**) from background regions in different reconstructions (n=15 from three cells for each sample). Yellow arrowheads in (**c**) and (**e**) indicate the inaccurate reconstructions of pure DL-based methods. Red arrowheads in (**e**) highlight the artifacts of physical-model-based methods. (**h-i**) SSIM of actin filaments (**h**) and ER tubules (**i**) in different reconstructions (n=150 and 15, respectively). (**j**) Resolutions of different reconstructions of actin filaments in **a-c** (n=14 from three cells). Scale bars: (**a**) and (**d**) 1 μm; (**b**) top, (**c**) and (**e**) 0.5 μm. (**b**) bottom, axial: 0.2 arbitrary units (a.u.); lateral: 0.1 μm.

Finally, we benchmarked the performance of TDV-SIM in resolving mitochondrial cristae dynamics for a prolonged time in live cells (**Fig. 3a**), which constituted an unresolved challenge for current SIM microscopy^15^. During the 20 s recording, the fluorescence intensity of Mito-Tracker decreased by ∼30% due to photobleaching (**Fig. 3b**). In the beginning, model-based methods could reconstruct high-quality intricate mitochondrial cristae, which were gradually corrupted with artifacts gradually due to photobleaching (**Fig. 3d-f**). In contrast, although pure DL-based methods consistently generated fewer artifacts during the imaging period, they could not predict most cristae structures in the first place (**Fig. 3c**,**e**,**f**). Outperforming all other methods, TDV-SIM obtained sharp mitochondrial cristae structures with fewer artifacts and high SSIM with the GT, which persisted even under photobleaching conditions (**Fig. 3c-f**).

**Fig. 3.**
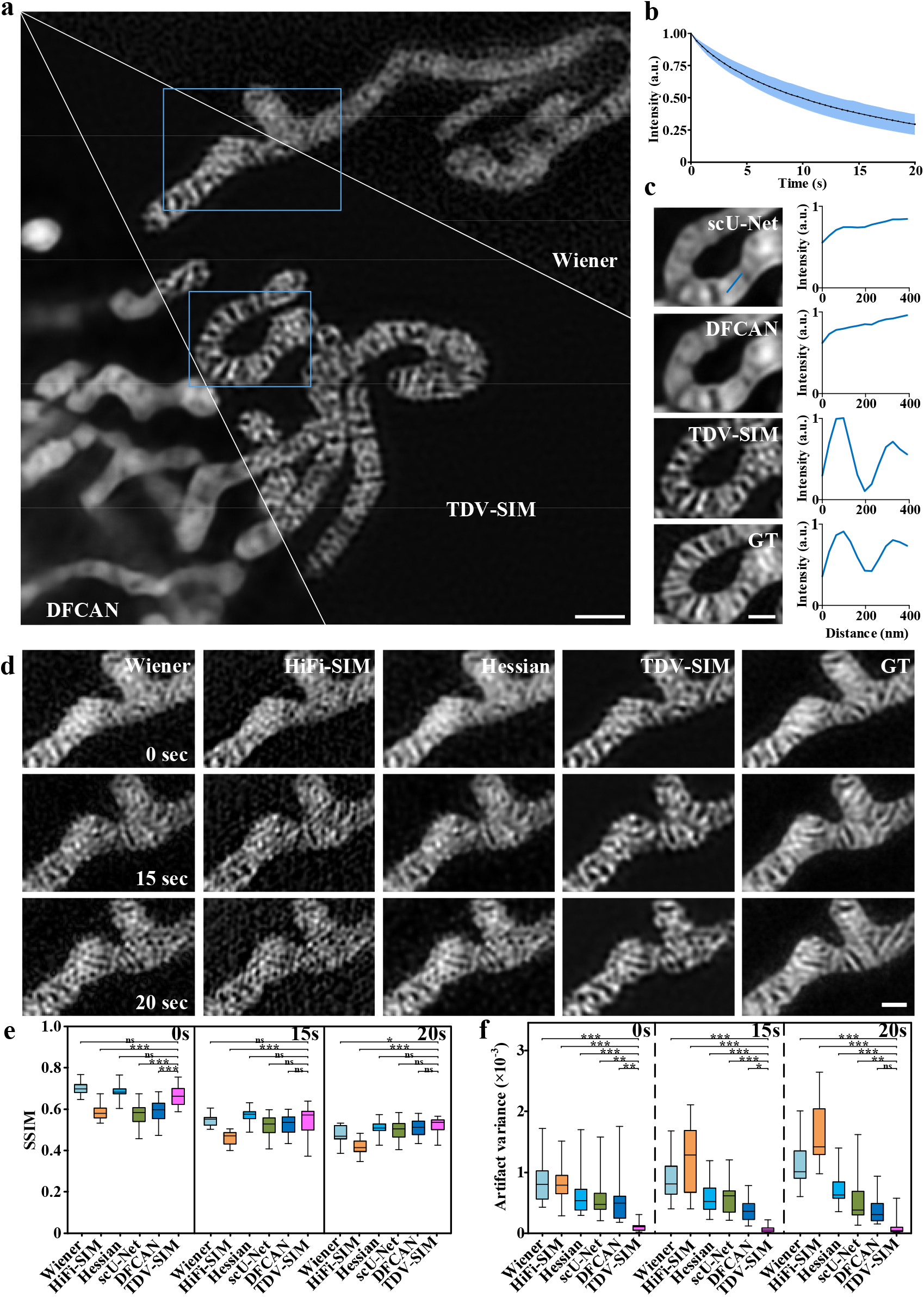
TDV-SIM enables accurate reconstruction of intricate and dynamic mitochondrial cristae structures in live cells after prolonged bleaching. (**a**) Mitochondria under the SR-SIM. (**b**) Time-dependent bleaching in fluorescence intensities of mitochondria. (**c**) Magnified views of the smaller boxed region in (**a**) reconstructed by scU-Net, DFCAN, and TDV-SIM and the corresponding GT image at 0 s. Profiles along the blue line are on the right. (**d**) Magnified views of the larger boxed region in (**a**) reconstructed by Wiener deconvolution, HiFi-SIM, Hessian-SIM, and TDV-SIM and the corresponding GT image at 0 s, 15 s, and 20 s. (**e**) The SSIMs of regions enclosed mitochondria from different reconstructions compared to GT images at 0 s, 15 s, and 20 s (n=15). (**f**) Artifact variances of the background regions in different reconstructions at 0 s, 15 s, and 20 s (n=15). Scale bars: (**a**) 1 μm; (**c**) and (**d**) 0.5 μm.

In summary, initiating from a hybrid angle, TDV-SIM presents a novel solution for high-resolution and high-fidelity SR-SIM reconstruction from low SNR raw data. Compared with pure DL methods, incorporating physical constraints about the image formation process is critical for TDV-SIM to maintain resolution and correctly predict irregular and complicated structures in constant changes. On the other hand, it also retains the benefit of DL methods in suppressing artifacts even under conditions with scarce photons available. Thus, TDV-SIM can obtain high-quality data during prolonged time-lapse imaging and will be crucial for SR imaging subcellular structure dynamics in live cells. Overall, combining the physical model with DL may represent an alternative and transferable way to reconstruct high-fidelity fluorescence images of different microscopes.

## Methods

### Cell culture and labeling

COS-7 cells (ATCC, CRL-1651) were cultured in high-glucose DMEM (Gibco, 21063029) supplemented with 10% FBS (Gibco) and 1% 100□mM sodium pyruvate solution (Sigma-Aldrich, S8636) in an incubator at 37□°C with 5% CO_2_ until reaching ∼75% confluency.

To label mitochondria, COS-7 cells were incubated with 250□nM MitoTracker Green FM (Thermo Fisher Scientific, M7514) in an HBSS medium (Thermo Fisher Scientific, 14025076) containing Ca^2+^ and Mg^2+^ at 37□°C for 15□min, followed by washing for three times before conducting 2D SIM imaging. To label actin, COS-7 cells were transfected with Lifeact–EGFP. According to the manufacturer’s instructions, the transfections were executed using Lipofectamine 2000 (Thermo Fisher Scientific, 11668019). After transfection, the cells were plated on precoated coverslips. Live cells were imaged in a complete cell culture medium containing no phenol red in a 37□°C live-cell imaging system. To label ER, COS-7 cells were transfected with EGFP-KDEL. According to the manufacturer’s instructions, the transfections were executed using Lipofectamine 3000 (Thermo Fisher Scientific, L3000015). After transfection, the cells were cultured for 20-28 h before the experiments. Live cells were imaged in a complete cell culture medium containing no phenol red in a 37□°C live-cell imaging system. The cells were tested for mycoplasma contamination before use.

### Image acquisition, preprocessing, and training

The same SIM settings in Hessian-SIM^8^ were used. To obtain low SNR raw images and the corresponding GT images for training the neural network, we imaged the specimen with SIM. We recorded 20 images for each illumination pattern and then changed the phase and orientation of the pattern. We repeated the cycle nine times, corresponding to three orientations multiplied by three phases, thus obtaining 180 raw images. Then we divided the raw images into 20 groups, with each group containing nine illumination patterns of three phases and three orientations. After removing the fluorescent background, we can obtain 20 SR images with artifacts using Wiener deconvolution. Finally, we mimicked the artifact-free ground truth (GT) by averaging the 20 SR images.

We imaged approximately 20 cells, and the images were preprocessed to obtain pairs of raw data and GT images at each time point. Next, we divided such image pairs into a training set, validation set, and test set; then, we applied random cropping, quarter rotating, and horizontal/vertical flipping to further enrich the training dataset. We trained the TDV-SIM using an Adam optimizer, with the learning rate set to 10^−4^. For actin, we adopted the mean square error (MSE) loss function:

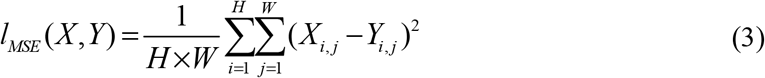

where W and H represent the image width and height, respectively. For mitochondria and ER, a combination of the MSE loss and the structural similarity index (SSIM) loss was used:

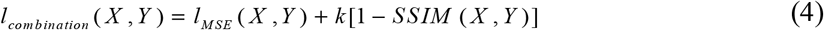

where *k* is a scalar weight that balances the relative contributions of SSIM and MSE losses and is set to 0.1 throughout this paper.

### Calculation of assessment metrics

To avoid the influence of different methods on the dynamic range of the inferred SR images, we first normalize the SR images:

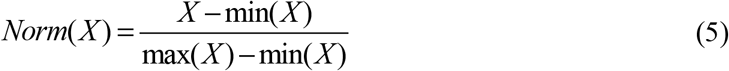

We used the PSNR, SSIM, and normalized root mean square error (NRMSE) to evaluate the similarity between the reconstructed image and GT. They were calculated as follows:

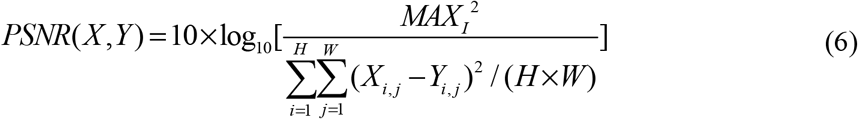

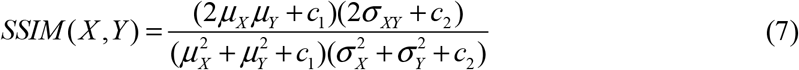

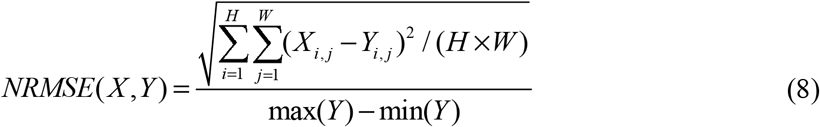

where *W* and *H* represent the image width and height, respectively. *X* and *Y* represent the reconstruction result and the GT image, respectively. *MAX*_*I*_ is the maximum possible pixel value of the image and equals to 2^*B*^-1 when the image is represented with linear pulse-code modulation of *B* bits (For example, *MAX*_*I*_ equals 255 for 8 bit image). μ*X* and μ*Y* represent the averages of *X* and *Y*, σ*X* and σ*Y* represent the variances of *X* and *Y*, and σ*XY* represents the covariance of *X*, and *Y. c*_*1*_ and *c*_*2*_ are small positive constants that stabilize each term; *c*_1_ = (0.01 *L*)^2^, *c*_2_ = (0.03 *L*)^2^, where *L* is the dynamic range of the pixel values.

Artifacts often emerged in regions of minor signals, such as the meshed region within actin filaments. Therefore, benchmarked against the GT, we selected these regions to calculate their variances.

## Data and code availability

All the data and code supporting this study’s findings are available from the corresponding author upon reasonable request.

## Acknowledgments

We acknowledge support by grants from the National Natural Science Foundation of China (92054301, 81925022, 92150301, 32170691, 62103071, and 31901061), the Beijing Natural Science Foundation (Z20J00059), the National Science and Technology Major Project Program (2021YFA1100201), the Lingang Laboratory (LG-QS-202206-06), Clinical Medicine Plus X - Young Scholars Project, Peking University, the Fundamental Research Funds for the Central Universities, the Natural Science Foundation of Chongqing (cstc2021jcyj-msxmX0526), the Science and Technology Research Program of Chongqing Municipal Education Commission (KJQN202100630), the Strategic Priority Research Program of Chinese Academy of Sciences (XDA16021200) and the High-Performance Computing Platform of Peking University.

## Author contributions

J.F. and J.W. carried out the experiments. J.W. and B.Z. performed the data analysis. J.W. and B.Z. drafted the manuscript. X.H. prepared and conducted the cell experiments. L.C. and X.H. conceived and designed the study and revised the manuscript. All authors read and approved the manuscript.

