## supplementary note 1~3, supplementary figure 1~3, video for "A hybrid reconstruction of the physical model with the deep-learning that improves structured illumination microscopy": hybridSIM supplementary v23.docx

**Supplementary Figures**

Supplementary Fig. 1| **TDV-SIM can restore the intricate structure of low SNR**. (**a**) The entire pipeline of simulation dataset generation. (**b**) The SR images reconstructed by different reconstruction algorithms. (**c**) The pipeline of segmenting the foreground and background. (**d**) Statistical comparison of different reconstruction algorithms in terms of SSIM, NRMSE, and artifact variance. (**e**) The SSIM and NRMSE of the foreground region in the SR images reconstructed by different algorithms in terms of SSIM and NRMSE.

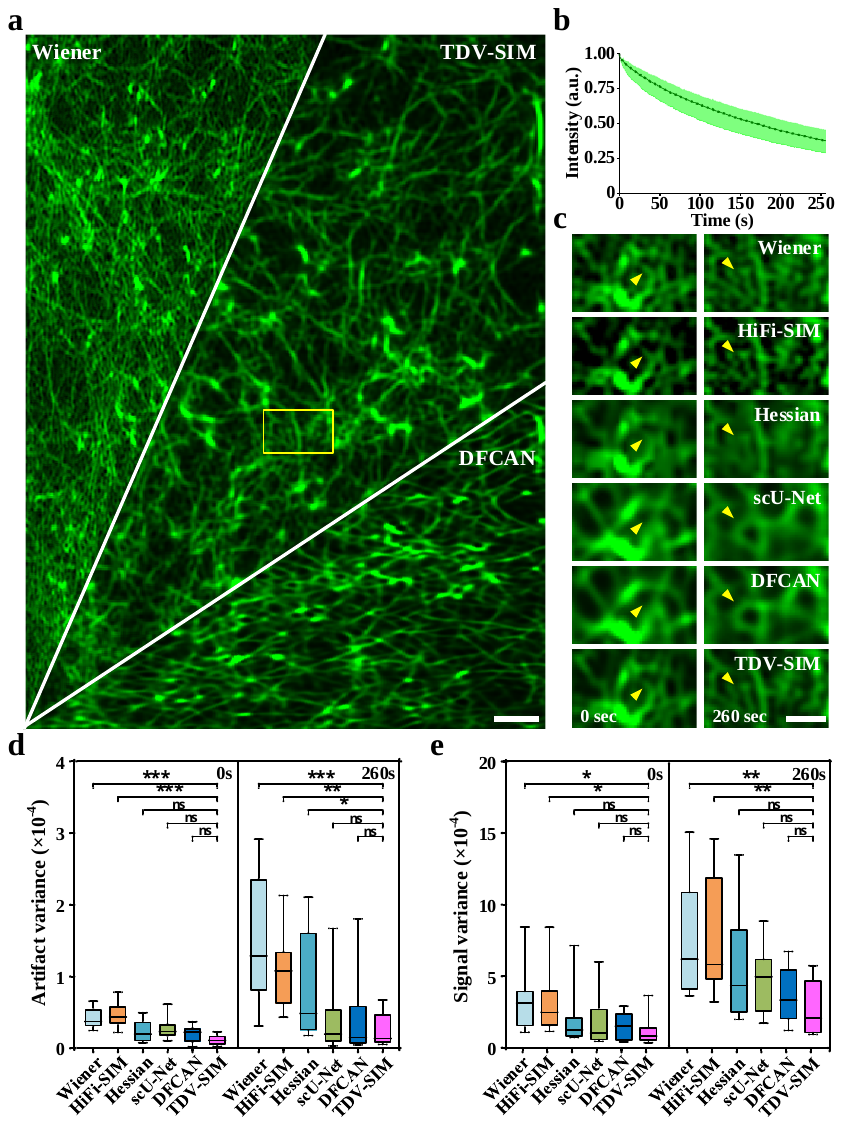

Supplementary Fig. 2| **TDV-SIM can restore regular structures of different SNRs without GT.** (**a**) A representative SR image of Lifeact-EGFP labeled actin in a live COS-7 cell. **(b)** Time-dependent fluorescence bleaching of Lifeact-EGFP. (**c**) Magnified views of the boxed region in (a) reconstructed via Wiener deconvolution, HiFi-SIM, Hessian-SIM, scU-Net, DFCAN, and TDV-SIM at 0 s and 260 s. (**d**) Artifact variances of the meshed region within actin filaments with different algorithms at 0 s and 260 s (n=10). (**e**) Fluorescence signal variance along the actin filaments with different algorithms at 0 s and 260 s (n=10). Scale bars: (**a**) 1 μm; (**c**) 0.5 μm.

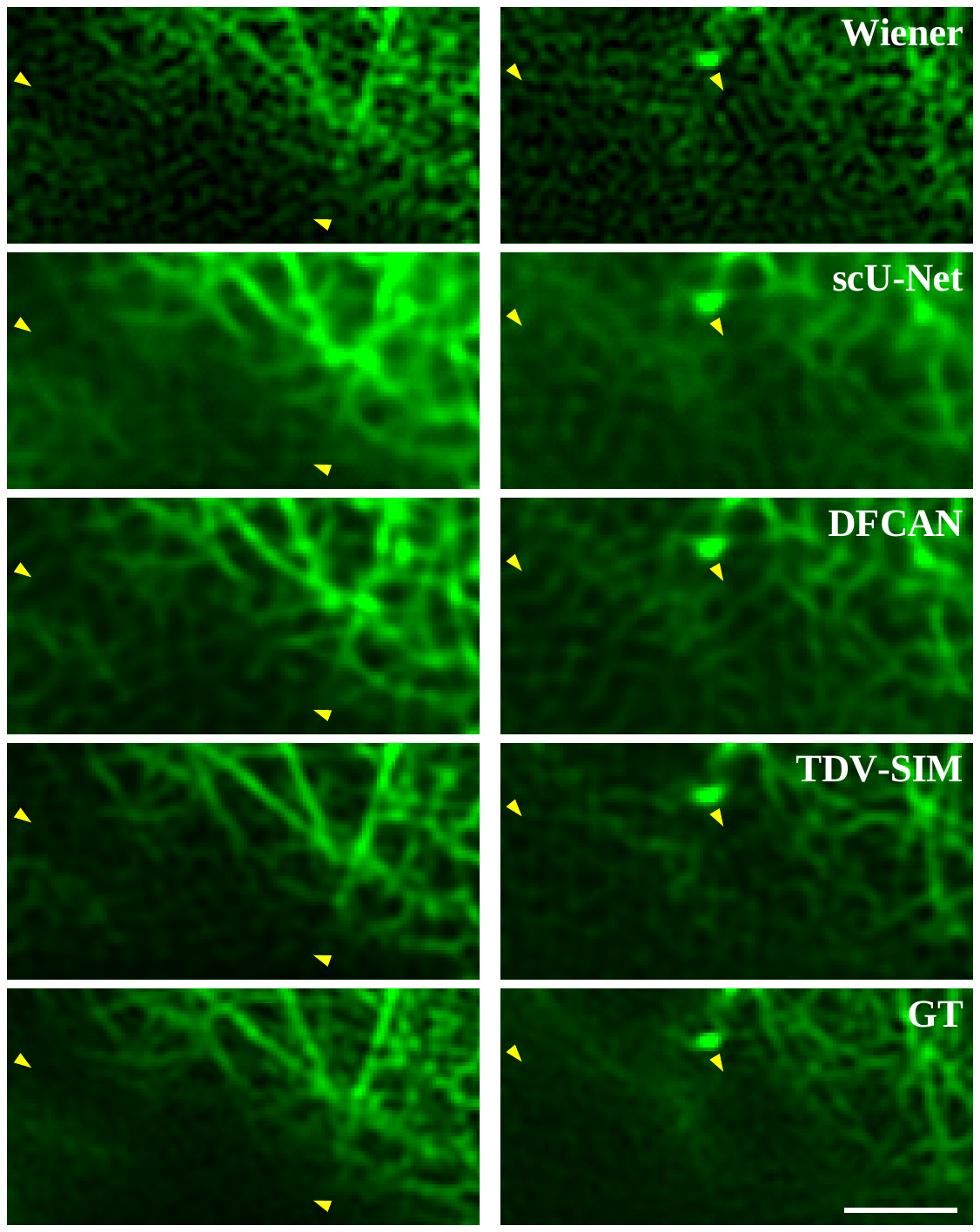

Supplementary Fig. 3 | **Pure DL-based methods infer actin filaments incorrectly at regions with extremely low fluorescence intensity.** Yellow arrowheads indicate the inaccurate reconstructions of pure DL-based methods. Scale bars: 1 μm.

**Supplementary Notes**

**Supplementary Note 1 | Fidelity and regularization term reconstruction in TDV-SIM**

When imaged by SIM, the sample is excited by sinusoidal illumination patterns with different orientations and phases. Raw images contain both low and high-frequency information about the sample, and the high-frequency information is scaled and downshifted into the optical transfer function (OTF) of the imaging system^1^. To reconstruct an SR image, the high-frequency information must be separated from the raw data before being assembled with the low-frequency information^2^. Here, the SIM reconstruction is formulated as an optimization problem:

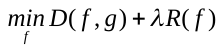

where *f* is the artifact-free SIM image to be estimated, g is the inverse Fourier transform of the assembled high and low-frequency bands, *D*(*f*, *g*) is the fidelity term based on the physics model of SIM, *R*(*f*) is the DL-based TDV regularization term^3^, and *λ* is the weight parameter of the regularization term.

To construct the fidelity term, we extract SR components that have been shifted into the OTF due to pattern illumination as follows:

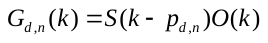

where *d* and *n* represent the pattern orientations and the orders of the bands, *p_d,n_* represents the pattern wave vector, *O*(*k*) represents the OTF, and *S*(*k*) represents the Fourier transform of the spatial distribution *s*(*r*) of the fluorophores in the specimen. In the spatial domain, **Eq.** can be rewritten as

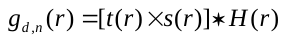

where *H*(*r*) is the point spread function (PSF) of the imaging system and *t*(*r*) is a phase factor that shifts the spectrum of *s*(*r*) in the Fourier domain:

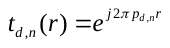

Since *g* can be obtained from **Eq.** , we use the *l_2_* norm of the difference between frequency bands extracted from the raw data and SR images as the likelihood term:

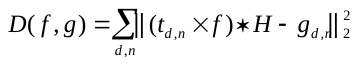

To simplify the calculation, we convert **Eq.** to the following form:

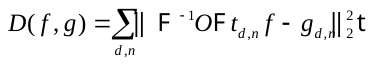

where  and ^-1^ represent the Fourier and inverse Fourier transform, respectively. Thus, the gradient of the likelihood term can be calculated as follows:

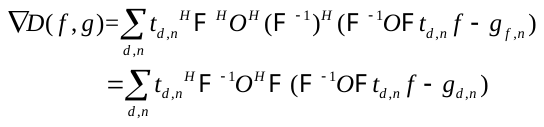

where the superscript *H* means conjugate transpose.

To construct the regularization term, we use TDV defined as the total sum of the pixel-wise deep variation:

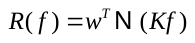

where *K* is a learned convolution kernel with zero-mean constraint,  is a multiscale convolutional neural network, and *w* is a learned weight vector. The exact form of TDV is depicted in **Fig. 1b**. The multiscale convolutional neural network (CNN) comprises three U-Net type architectures. Each U-Net type architecture consists of five micro-blocks with a residual structure on three scales. Next, to compute the gradient of the regular term, ∇*R*(*f*), we inverted its architecture, transformed the activation function into its gradient, and turned the convolution layer into the transpose convolution layer with the same kernel (**Fig. 1b**).

**Supplementary Note 2 | TDV-SIM excels in restoring structures of low SNR**

Because the ground truth (GT) images of the actual images were obtained by averaging the results of Wiener deconvolution and were not the actual fluorescence distribution, this may affect the final evaluations of SR reconstruction qualities by different methods. Therefore, we carried out a simulation experiment (**Fig. S1**).

We processed the high-resolution (HR) images of the DIV2K^4^ dataset to obtain the pairs of simulated raw SIM images and GT images (**Fig. S1a**). We randomly cropped a square area from the RGB image, transformed it into a grayscale image, extracted its edges, and then multiplied the edge image and grayscale image to simulate the fluorescence distribution in the biological sample. Next, we multiplied the fluorescence distribution image with the illumination patterns of three directions and three phases to obtain the simulated SIM raw images. Then, the imaging process was simulated by multiplying the optical transfer function (OTF) of the wide-field (WF) microscope in the frequency domain. The acquisition process of the camera was simulated by downsampling. Finally, we add Gaussian noise with a mean of 0 and variance of 0.005 to the images. To obtain the simulated GT image, we multiplied the simulated fluorescence distribution image in the frequency domain by the equivalent OTF of SIM, which was obtained by moving the OTF of the WF microscope in the frequency domain according to the illumination parameters, then binarization, and multiplying by the apodization function, where the apodization function is defined as:

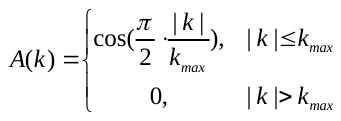

where *k* represents the frequency coordinate, *k_max_* is the upper limiting frequency set, and the OTF radius. We re-divided the dataset into the training, validation, and test sets in a 6:1:2 ratio.

We compared the performance of TDV-SIM with other reconstruction algorithms, including Wiener deconvolution^5,6^, HiFi-SIM^7^, Hessian-SIM^2^, scU-Net^8^, and DFCAN^9^ on the test sets (**Fig. S1b**). To evaluate the similarity of these SR images to GT, we calculated their structural similarity index (SSIM) and normalized root mean square error (NRMSE). In addition, to evaluate the artifacts, we performed Gaussian filtering on the GT images, segmented the foreground and background by thresholding, and then calculated the variance of the artifacts in the background region (**Fig. S1c,d**). Wiener deconvolution, HiFi-SIM, and Hessian-SIM produced more artifacts in the SR images, resulting in lower SSIM, higher NRMSE, and higher artifact variance. The artifact levels of images reconstructed by scU-Net and DFCAN are significantly lower and close to TDV-SIM. However, when we focus on the image's foreground and calculate the SSIM and NRMSE of the foreground region, we find that TDV-SIM has obvious advantages compared with other methods (**Fig. S1e**). These results suggest that TDV-SIM has an advantage in recovering intricate structures from low SNR raw data.

**Supplementary Note 3 | TDV-SIM excels in restoring regular structures of different SNRs without GT**

In practical applications, GT is unknown, and we used Wiener deconvolution as the reference for providing the real structure information. We compared the performance of TDV-SIM with other reconstruction algorithms, including Wiener deconvolution, HiFi-SIM, Hessian-SIM, scU-Net, and DFCAN on actin (**Fig. S2a**). Actin filaments were labeled with Lifeact-EGFP and captured with a 2.7 ms exposure. During the 260 s recording, the fluorescence intensity of Lifeact-EGFP decreased by ~63% due to photobleaching (**Fig. S2b**). Compared with the conventional Wiener deconvolution, reconstruction algorithms based on the physical inversion model (HiFi-SIM, Hessian-SIM) can faithfully preserve the sample structure information but produce significant artifacts in the background regions (**Fig. 2b** and red arrows in **Fig. 2e**). In contrast, reconstruction algorithms based on DL (DFCAN, scU-Net) can suppress the artifacts effectively but predict structures deviating from the Wiener deconvolution (yellow arrows in **Fig. S2c**). As a representative method combining the physical model with the DL-based regularizer, TDV-SIM can preserve the sample structure information faithfully and suppress the artifacts effectively at the same time (**Fig. S2c**). In line with the visual inspection results, TDV-SIM restoration demonstrated the lowest artifact variance, which was more significant than the physical model-based methods, and similar to that of the DL-based methods (**Fig. S2d**). In addition, TDV-SIM restoration demonstrated the lowest fluorescence signal variance along the actin filaments, which was more significant than the Wiener deconvolution and HiFi-SIM, and similar to that of Hessian-SIM, scU-Net, and DFCAN (**Fig. S2e**). Collectively, these data indicate that TDV-SIM outperforms other algorithms in restoring high fidelity and low artifact SR images from raw data of different SNRs without GT.
